# Ecosystem services in connected catchment to coast ecosystems: monitoring to detect emerging trends

**DOI:** 10.1101/2021.07.19.453015

**Authors:** Christopher J. Brown, Chantal Saint Ange, Rod M. Connolly, Syezlin Hasan, Sue Jackson, Joseph M. McMahon, James C. R. Smart

**Author notes:** Corresponding author 170 Kessels Road Nathan, Queensland, Australia, 4111.

## Abstract

There is an increasing need for long-term monitoring of ecosystems and their services to inform on-ground management and policy direction. The supply of many ecosystem services relies on connections that span multiple ecosystems. Monitoring the underlying condition of interconnected ecosystems, using established indicators, is therefore required to track effectiveness of past interventions and, ideally, identify impending change. Here we conduct performance testing of ecological indicators for a catchment-to-coast system with the aim of identifying the time-scales over which they respond to change. We chose a case-study of a coastal fishery in Northern Australia that exhibits strong catchment-to-coast connectivity, has long-term available data and is under threat from water resource development. We developed a novel approach to performance testing. Our model drew on state-space modelling to capture ecological dynamics, and structural equation modelling to capture covariation in indictors timeseries. We first quantified covariation among three established ecological indicators: pasture biomass, vegetation greenness and barramundi catch per unit effort. Covariation in the indicators was driven by river flow, with higher values of all indicators occurring in years with greater river flow. We then defined reference bounds for each indicator that accounted for natural variation in river flow. We predicted the emergence times for each indicator, as the time taken for each indicator to emerge from the background of natural variation. Emergence times quantified at 80% and higher confidence levels were >10 years in all cases. Past trends and current status of ecosystem service flows are often used by decision makers to directly inform near-term actions, particularly provisioning services (such as barramundi catch) due to their important contribution to regional economies. We found that the ecological indicators should be used to assess historical performance over decadal timespans, but not as short-term indicators of recent change. More generally, we offer an approach to performance testing of indicators. This approach could be useful for quantifying time-scales of ecosystem response in other systems where cross-ecosystem connections are important.

## Introduction

The supply of ecosystem services can depend on ecological processes that operate across multiple ecosystems (Barbier et al. 2011; Lowe et al. 2022). Interconnections among ecosystems can occur through the movement of plant propagules, migration of animals and the transport of physical matter in air and water. In particular, the flow of water from catchments to coast creates many challenges for the management of human activities that may impact upon the supply of ecosystem services. For example, the capture of water in dams for agriculture, power or municipal uses upstream interferes with the supply of ecosystem services all the way to the ocean, potentially affecting ecosystem inputs to agriculture on floodplains, the productivity of estuarine and coastal marine fisheries (Gillanders et al. 2011) and the capture of carbon in estuarine habitats (Macreadie et al. 2017). Therefore, monitoring of multiple ecological indicators that together span interconnected systems is important to characterize the processes that support ecosystem services in catchments and inform the management of human activities. This monitoring of ecological processes plays multiple roles in the decision-making. First, it informs decision makers about trends in downstream ecosystem processes so they can respond proactively to address causal changes in hydrology upstream. Second, it helps to detect change in ecosystems and attribution to management actions. Finally, monitoring can hold decision makers accountable for actions that affect the integrity of ecosystems downstream (Stevenson et al. 2021).

A challenge for monitoring that spans a whole catchment is to attribute trends in ecological processes to changes in water flow, because the causes of changes in water flow are separated in time and space from their impacts (Alvarez-Romero et al. 2011; Brown et al. 2019). Interannual variation in flow linked to long-term climate cycles (e.g. El Nino southern oscillation) is a common property of many systems, and this variation also contributes to masking of long-term trends caused by climate change or human interventions. Monitoring design, such as ‘before/after, control/impact’ approaches can be used to resolve the attribution problem (Underwood 1992). However, in whole of catchment monitoring programs, commonly no controls exist; monitoring programs are not designed with a specific intervention such as water extraction for irrigation as the focus. In such cases we need to know how long underlying trends in hydrology/flow will take to manifest as trends in the ecological indicators.

The testing of indicator performance, where the ability to detect trends in indicators is explored through simulation models, is a popular method in fisheries science for evaluating indicators (Fulton et al. 2005; Blanchard et al. 2014; Nickols et al. 2019), but it is only just beginning to be applied to ecosystem indictors (Collen & Nicholson 2014; Watermeyer et al. 2021). Performance testing involves constructing a system model that represents the fundamental system dynamics, and then running simulations that represent the stages of underlying trends, data collection and interpretation of data by decision-makers (Fulton et al. 2005). Simulations are run multiple times so that we can define error bounds on the probability of a trend of a given magnitude being detected with a given level of confidence. This testing can be evaluated through univariate indicators, such as binary indicators of whether a threshold value is reached, or via power analyses that consider the sensitivity and specificity of indicators for different decision thresholds (i.e. risk of missed detections versus false alarms - Mapstone 1995). The development of performance testing methods in fisheries science and biodiversity monitoring has run parallel to the development of the emergence time concept in climate studies (Hawkins & Sutton 2012). Climate models can be used to quantify when human induced trends in climate caused by greenhouse gas emissions emerge from background variation in climate. Thus, models of trends underpin the attribution of climate trends to human causes and provide an easily communicable statistic about lags between human actions and detection of changes in the system under study. The emergence time concept could also be applied to the testing of indicators for interconnected catchment ecosystems, where high interannual variation in river flows may mask long-term trends in multiple downstream ecological processes. However, development of such performance testing is often held back by the requirement to develop complex system models. Developing system models for catchment to coast systems is a particularly complex task because the modelling needs to span multiple disciplines (Brown et al. 2019). As such, catchment to coast systems models are rare. Simpler approaches to performance testing would facilitate more widespread testing of indicators, and therefore inform how managers can use indicators.

Here we develop an approach to quantifying emergence time of multivariate catchment processes. We use a case-study of a riverine-estuarine-coastal marine system in the Gulf of Carpentaria in tropical northern Australia. The Mitchell catchment is earmarked for potential water resource development, which may provide agricultural benefits, but will also disrupt freshwater flows (CSIRO 2018) and impact the downstream ecosystems and fisheries that depend on them (Turschwell et al. 2019; Broadley et al. 2020). Similarly, climate change is causing long-term shifts in weather patterns and flows in the region (Karim et al. 2015). We first selected commonly used monitoring indicators for the interconnected ecosystems of the Mitchell catchment. We then sought to fit a process model that quantified the relationship among indicators and that allowed the effect of uncertainty within those indicators to be assessed (Aim 1). From this model we inferred natural bounds of variation for each indicator (Aim 2). We finally quantified the sensitivity of indicators to long-term trends in water flow via their emergence time from background variation (Aim 3). The emergence time for each indicator represents when the impacts of change will become apparent to human observers (Hawkins & Sutton 2012). Overall, our approach provides a method for performance testing of multiple indicators. Our approach could be used to support development of indicator sets for attribution of trends in human impacts, ecological condition accounts (Czúcz et al. 2021) and monitoring of human impacts.

## Methods

### Indicators for the Mitchell catchment

The Mitchell catchment covers an area of over 72,000 km^2^ and discharges into the Gulf of Carpentaria. This catchment is subject to highly seasonal flow regimes. During the dry season many of the ephemeral rivers become disconnected water holes and in the wet season a large area becomes inundated. There is high inter-annual variability in rainfall, which drives inter-annual variation in flow and floodplain productivity (Ndehedehe et al. 2021). We therefore selected variables and indicators for the catchment with the aim of capturing variation in ecological states related to river flow.

We focused on indicators of capacity to supply ecosystem services that included (i) fish biomass that supports barramundi (*Lates calcarifer*) catch in a commercial fishery and resource harvesting of barramundi by Indigenous Australians, and (ii) pasture biomass that supports commercial livestock production. In particular, the catadromous barramundi life-cycle exemplifies cross-system dependence (Barramundi migrate from freshwater to estuaries to spawn, Milton and Chenery 2005). Variability in barramundi biomass may be correlated to variability in pasture growth on flood plains which supports livestock production, because both are dependent on regional cycles in river flow.

Time-series data (Table S1) were collected for the ecosystem service capacity indicators, and additional indicators where long-term data were available. Stream flow, a hydrological indicator, was included and sourced from the Northern Australia Water Resource Assessment (NAWRA).

Yearly data on barramundi (*Lates calcarifer*) catch and fishing effort were used to derive an indicator of the barramundi population state. Data were sourced from the Queensland Fisheries ‘QFish’ data cube (State of Queensland Department of Agriculture Fisheries and Forestry 2018) for the Mitchell region. Barramundi are an important fishery species in northern Australia, and they are caught in commercial, Indigenous, recreational and charter fisheries. They also play an important cultural role within the ecosystem (Jackson et al. 2012). Barramundi spawn in estuaries, with juveniles then moving upstream to freshwater habitats, returning to saltwater as adults (Robins et al. 2005).

Seasonal floodplains provide connection to off-stream freshwater habitats as well as highly productive, short-term habitats. Individuals residing in upstream habitats return to the estuary and saltwater to spawn as adults. Barramundi are impacted by flow regimes through changes in connectivity throughout the catchment which benefit growth and survival (Robins et al. 2005; Balston 2009; Roberts et al. 2019; Leahy & Robins 2021). Barramundi opportunistically exploit multiple aquatic habitats at different stages of their life cycle. Therefore, if connectivity between these different ecosystem assets is diminished, this can impact their harvestable biomass in following seasons. Barramundi catch per unit effort (CPUE) is a suitable indicator of the interlinked aquatic ecosystems, because barramundi are a near-apex predator in the system so their production aggregates multiple trophic levels, and they depend on persistent freshwater in lagoons and waterholes to survive dry periods (Robins et al. 2005).

Normalized difference vegetation index (NDVI) is an indicator of vegetation greenness (Pettorelli et al. 2005) and was obtained from the Bureau of Meteorology (Table S1). Pasture biomass was included as an indicator of supply of forage for grazing. Pasture biomass was obtained as the predicted output from Queensland Government’s AussieGRASS model. AussieGRASS also uses stream flow as input data; however, since other variables (i.e. rainfall, soil typ, temperature and local grazing pressure) are also inputs to AussieGRASS (State of Queensland Department of Environment and Science, 2021), pasture biomass and stream flow are not highly correlated.

### Aim 1: Quantify interrelations among condition indicators

We combined two common modelling approaches to develop a model of indicator interrelationships (fig. 1). The first was a state-space model of barramundi catch per unit effort (CPUE) dynamics (Millar & Meyer 2000). State space models are widely applied for modelling fisheries dynamics, informing on reference points for fisheries management and for estimating environmental effects on fishery dynamics (Aeberhard et al. 2018), but have rarely been applied to model whole ecosystem change (Auger-Méthé et al. 2021). The state-space model was used to fit catches (a measure of ecosystem service flow), then estimate latent states for stock biomass (a measure of ecosystem capacity to deliver services) and environmental process variation (fig. 1).

**Figure 1.**
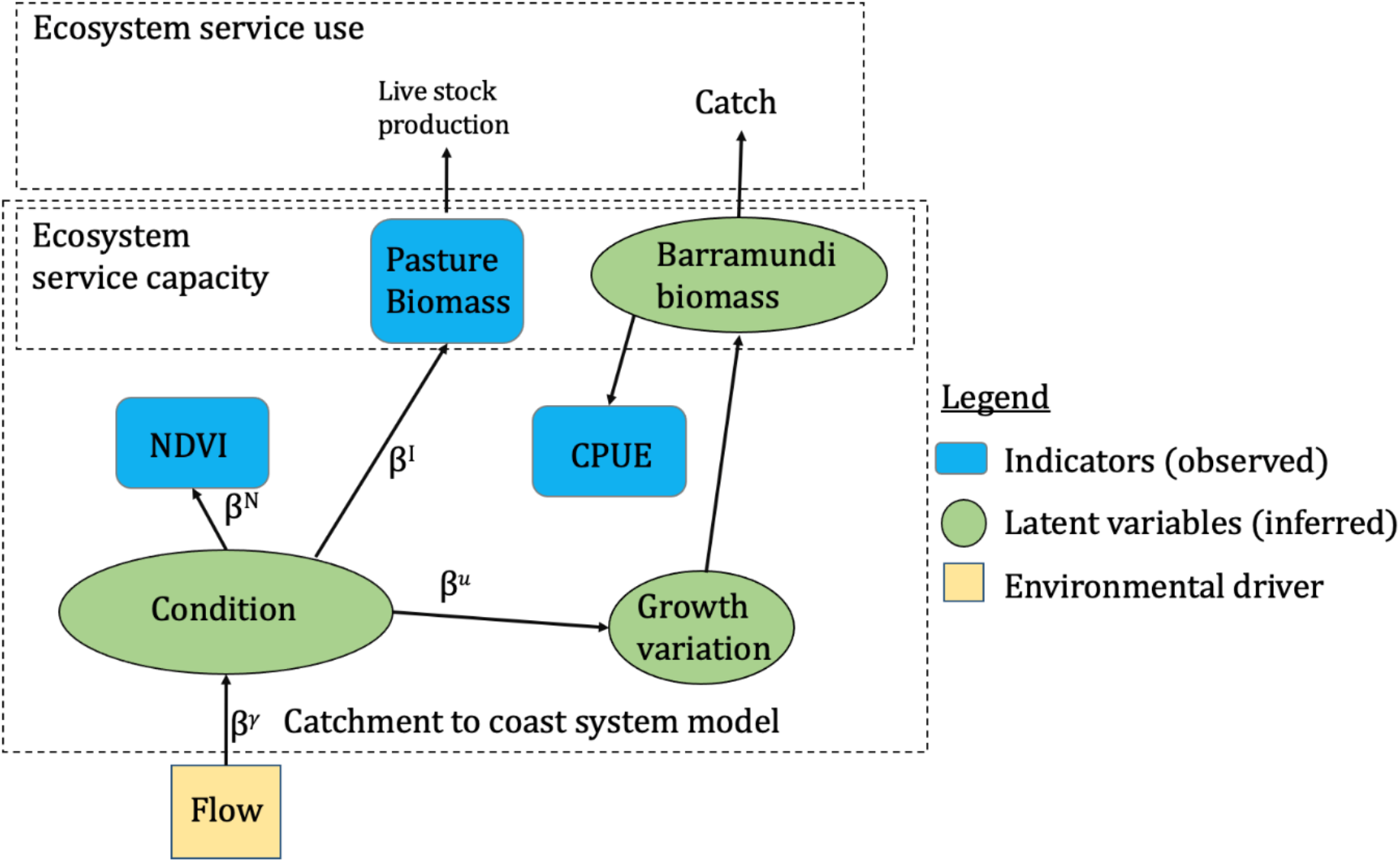
Overview of the approach showing the measured variables, the latent variables and the scope of the model.

The second modelling approach was a structural equation model that related the observed indicator variables to a latent (unobserved) state, which we termed ‘ecological condition index’ (fig. 1). Structural equation models are widely used to develop indicators, because they can model the correlations between observed indicators and a latent state that is hypothesized to be equivalent to the joint condition of different components of an ecosystem (Brown & Hamilton 2018). We linked the two modelling approaches by assuming that the observed indicators of NDVI, pasture biomass, and CPUE were all dependent on an unobserved ecosystem condition index. Ecosystem condition, in turn, was assumed to be driven by freshwater flow (fig 1).

The state space model had the form:

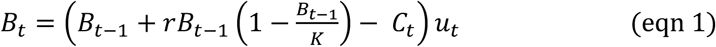

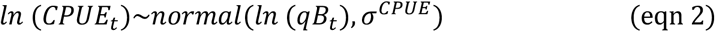

where *B_t_* was the latent estimate of biomass, r was the intrinsic growth parameter, *K* was the carrying capacity, *C_t_* was catches (assumed fixed), *q* is the catchability coefficient, *u_t_* were process errors and *σ^CPUE^* was the standard deviation for CPUE errors (we use superscripts to demarcate parameters relating to different variables, subscripts to demarcate indices). *B_1_* was assumed a fixed fraction of 20% of the estimate of *K* (Millar & Meyer 2000; Streipert et al. 2019). We tested the effects of the assumptions that *B_1_* was 20% of *K* and that *q* was constant over time in additional sensitivity analyses (Supplementary material).

The state space model was linked to the structural equation model through a dependence of the process errors on a latent variable we interpreted as the ecosystem condition index:

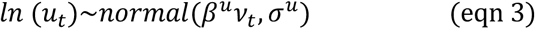

where *β^u^* was a regression coefficient and *v_t_* was the latent condition index. The process errors were assumed to be log-normally distributed so they had a multiplicative effect on biomass. This formulation is consistent with population theory because changes in survival have multiplicative effects on population size.

The state-space model assumed a one-year lag between flow and changes in barramundi biomass. Flow likely impacts barramundi catches and biomass at a number of different time-scales, as described above, but we assumed the single year impact for simplicity. To further verify this assumption, we confirmed that residuals from the modelled CPUE relationship had no autocorrelation. Future modelling could potentially separate the different types of flow effects on barramundi populations. But note this would require additional data types, because biomass, productivity and catchability parameters are close to being interchangeable in unstructured surplus production models, so they are difficult to estimate simultaneously.

NDVI and pasture biomass were also related directly to the condition index:

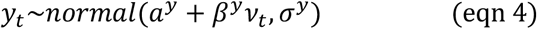

where *y_t_* refered to either NDVI or pasture biomass and *a^y^* is the intercept.

NDVI and pasture biomass were mean centred and standardized prior to analysis.

Finally, ecosystem condition was estimated as a regression on freshwater flow:

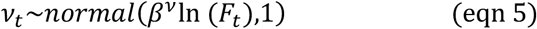

Where *F_t_* was freshwater flow and the standard deviation was arbitrarily set at 1, because *v_t_* is scale invariant. In our regression we used the log of flow, to reflect hypothesized multiplicative effects of flow on ecological condition.

The NDVI data started in 1992, so the condition index was estimated only from flow, CPUE and pasture biomass in 1990 and 1991. The model also interpolated the missing NDVI values. We presented results as marginal predictions for each data type and the latent states for barramundi biomass and ecological condition.

We obtained parameter and prediction probabilities within a Bayesian estimation framework. We set priors based on prior information for each parameter and prior expectations for the variance in NDVI and CPUE. It was also important to set weakly-informative priors to aid algorithm convergence and identification of parameters. In particular, the latent state *v_t_* and the *β* coefficients can ‘sign-switch’ (Hui et al. 2015). Prior choices and algorithm settings are explained in the Supplementary material. The model assumptions were checked by confirming independent normally distributed residuals, where residuals were defined as the observed values of CPUE, NDVI and pasture biomass minus their posterior predicted median value. Residual independence was confirmed by plotting the auto-correlation function with the R function ‘acf’ (R Development Core Team 2019). Further, we calculated the cross-correlation of residuals across indicators to confirm the model had captured all dependencies between indicator values. We further verified the model through a simulation study, focussing on the model’s ability to interpolate missing data (Supplementary material).

We determined the strength of the relationship between each indicator variable, ecological condition, and flow as the direct effect of condition on each indicator (simply the marginal posterior distributions for *β*^y^) and the total effect of flow (logged) on the condition index = *β^v^.β^y^*. For barramundi we calculated the direct and total effect sizes on surplus production, where surplus production was defined as the sum of change in biomass and catch (i.e. SP_*t*_ = *B_t_* – *B*_*t*–1_ + *C_t_*). We then plotted the total effect of a doubling of flow on each indicator and the direct effect of a one S.D. change in ecological condition on each indicator (see Supplemental methods). We also calculated one-tailed probabilities that the total effect was greater than zero. We note that given the prior choices (Table S2) the prior probability that *β^v^.β^y^* >0 was 0.5.

### Aim 2: Bounds of natural variation

We used the fitted model to define the bounds of natural variation for each indicator. The bounds were defined to represent the typical ranges of indicator values given historical variation in flow. The historical condition of flow was characterized by fitting an autoregressive moving average model to the historical flow series (1990-2015), then simulating 1000 random flow series from the fitted model (Supplementary material). To define bounds for the indicators we simulated their values for each of the 1000 flow series. For NDVI and pasture biomass, the bounds were then defined as the marginal probability quantiles integrating over measurement noise (also called predictive intervals) and the flow simulations. The bounds for barramundi CPUE also depended on biomass, so for simulations of barramundi we assumed a biomass fixed at the biomass that gives maximum sustainable yield (B_1_ = 50% of B_0_). This is a common reference point used in fishery management (Mace 1994). For the simulations of bounds, we also assumed that biomass was initially at the biomass that gives maximum sustainable yield, doing so avoided transient dynamics affecting the simulations.

### Aim 3: Emergence time for indicators

We then estimated each indicator’s emergence time, given an assumed long-term trend in flow. Climate change is causing long-term changes in flows in the region (Karim et al. 2015), which may affect interconnected catchment ecosystem services, including barramundi catches and pasture growth. Thus, our scenarios represent the ability to detect this long-term change under an assumed trend magnitude. We simulated flow series and added a linear decline in flow, as an example of a situation where environmental change or water resource development could put increasing constraints on ecosystem services. We simulated 1000 by 30-year time-series for each indicator and each trend in flow. We then defined emergence times as the time taken for the mean indicator value to cross the lower 5%, 20% and 40% probability quantiles. The different quantiles represent different degrees of confidence for emergence, the lower the quantile the greater the confidence the trend has emerged from background variation.

## Results

### Aim 1: Interrelationships among indicators

Verification of the model fits indicated that there was no autocorrelation in the residuals for the fitted values for any of the indicators (Fig S1). Overall, the R^2^ was highest for log CPUE (0.91), moderate for NDVI (0.43) and lowest for pasture biomass (0.09) (Fig S2). Sensitivity analyses in which we varied the assumptions regarding constant catchability, the initial biomass ratio and higher fishing effort indicated the main results were robust to alternative assumptions (Figs S2, S3 and S4). Consequently, below we present only the main results assuming stable catchability and an initial biomass ratio of 20%.

Model fits indicated an overall increase in CPUE from 1990 to 2012, and then a slight decrease from 2012-2015. NDVI and pasture biomass had no long-term trends, but varied year-to-year (Fig. 2). NDVI and barramundi surplus production were strongly related to river flow (Fig. 3A, pr(*β^v^.β^y^*.)>0 = 0.99 and 0.96 respectively), whereas pasture biomass had a somewhat weaker relationship with river flow (figure 3A, pr(*β^v^.β^y^*.)>0 = 0.93). Therefore, the ecological condition variable primarily captured the correlation between flow, NDVI and barramundi biomass change (Fig 2E, Fig 3B). Ecological condition showed peaks in 1991, 1998-99 and around 2010 that were driven by increased flow in those years. But overall ecological condition did not vary strongly from the long-term average (95% CIs generally overlap zero). Barramundi biomass was predicted to have increased from 0.2 to 0.5 as a fraction of B_0_ over 1990-2012, and to have declined to ^~^0.3 by 2015. The decline in 2012-2015 corresponded to a decline in ecological condition and a decline in flow.

**Figure 2.**
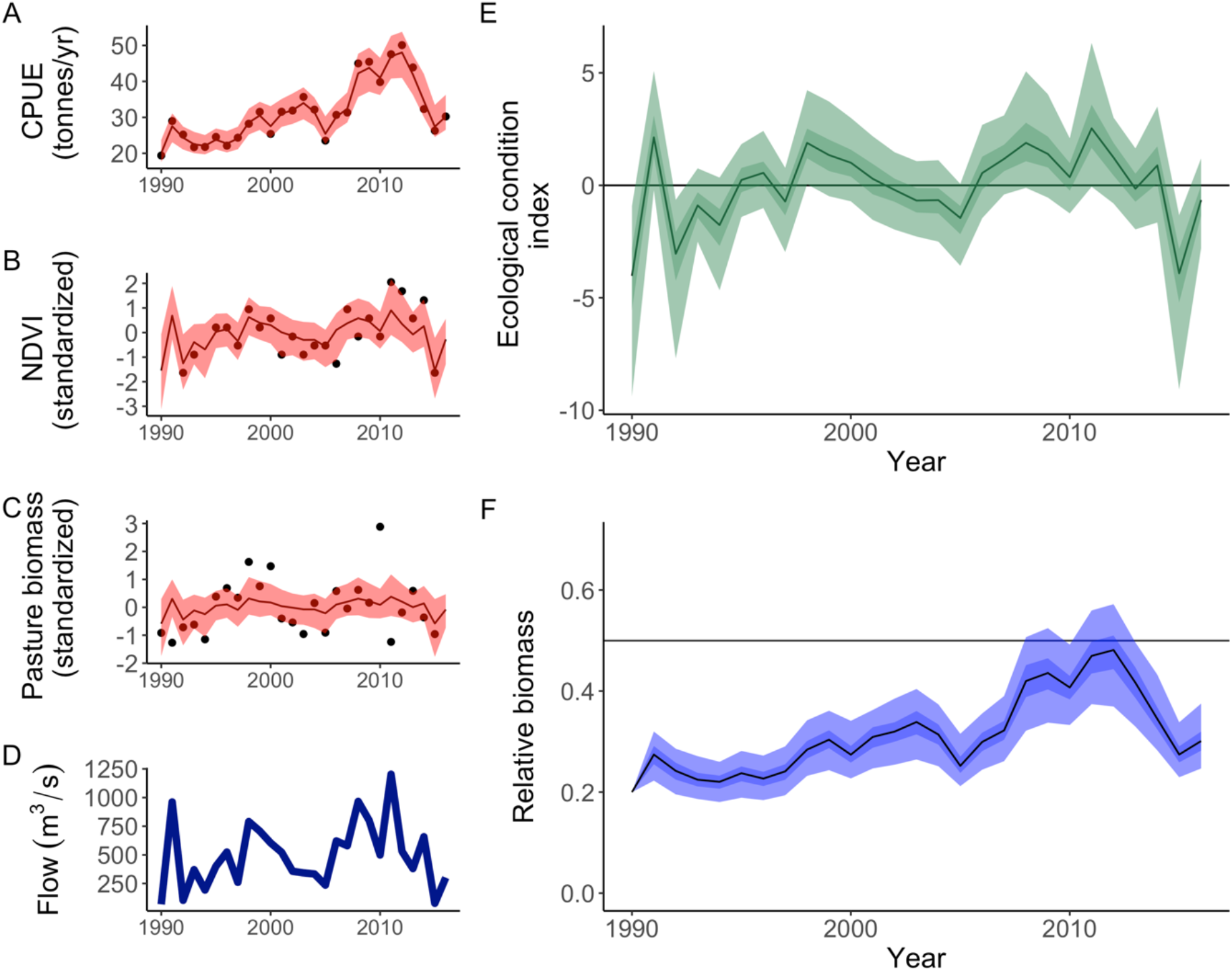
Model fits to data for CPUE (A), NDVI (B); pasture biomass (C); flow (D) and the estimated latent variables for ecological condition (variable *v_t_* in the model) (E) and barramundi biomass relative to its carrying capacity as estimated by the model (F). For model predictions, lines show median and shaded areas show 95% CIs. Darker areas in E-F indicate 50% CIs. The solid horizontal line in (E) indicates the mean value, in (F) it indicates BMSY of 50% carrying capacity.

**Figure 3.**
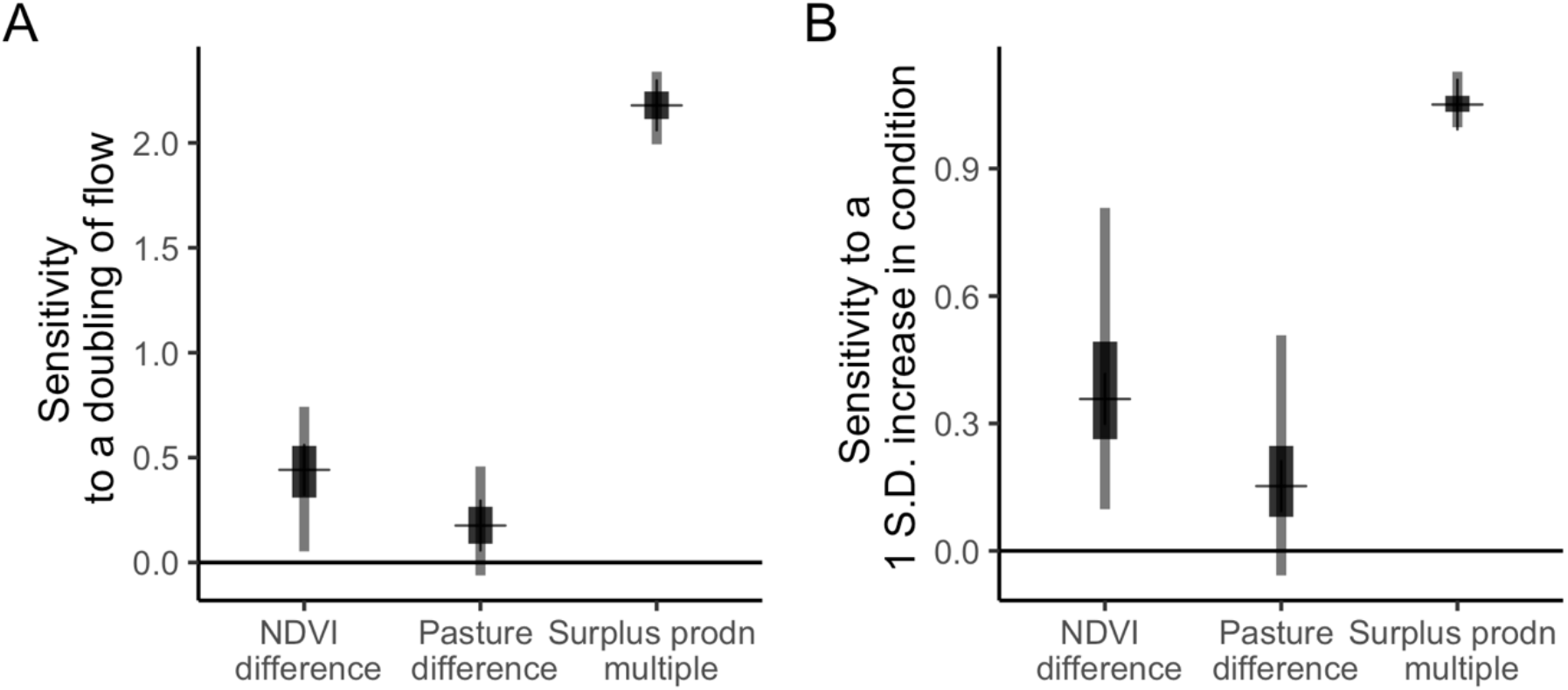
Strength of the relationships between each indicator and (A) a doubling of flow; and (B) a 1 S.D. increase in the ecological condition index. Note that NDVI and pasture are differences on a scale standardized to 1 S.D., whereas surplus production is on a multiplicative scale. Horizontal bars show median, thick solid lines show 75% probability quantiles and thin grey lines show 95% probability quantiles.

A simulation study where we varied the number of years of missing NDVI data suggested the model’s predictions for the variables and ecological condition were not biased by missing data, but the predictions for NDVI were less accurate with a greater amount of missing data (Fig. S5). Coverage by the 95% C.I.s was always >0.95 (Table S3). The parameter estimate for the effect of ecological condition on NDVI was most affected by missing data, and the parameter estimates for pasture biomass and barramundi biomass were least affected.

### Aim 2: Bounds of natural variation

The bounds for each indicator had broad intervals (Fig. 4), for instance, for CPUE the 80% probability interval ranged from 39-65 tonnes per fishing year, compared to a historical range of 19-50 tonnes per fishing year. This broad variation in the probability bounds was caused by the high variation in year-to-year flow.

**Figure 4.**
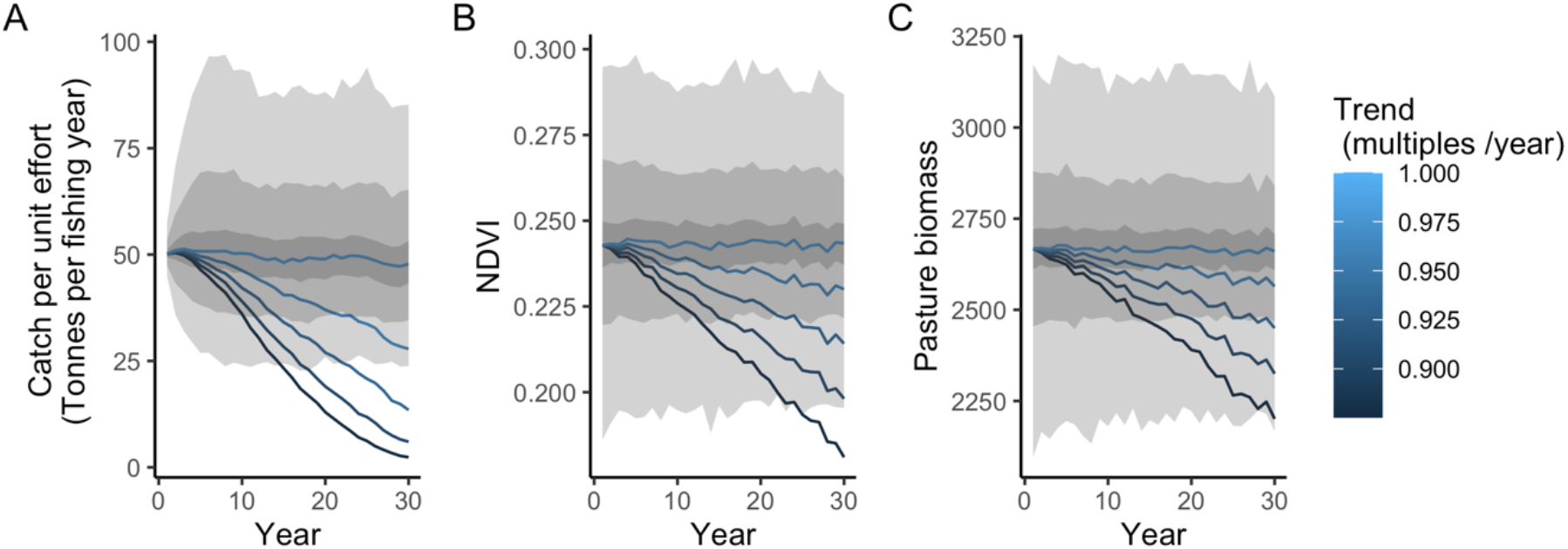
Bounds for catch per unit effort (A) NDVI (B) and pasture biomass (C). Grey zones show 40% (dark grey), 60% (medium grey) and 90% (light grey) probability intervals. Examples of trend lines shown in blue shades, as proportion of the mean.

### Aim 3: Emergence times

Emergence times were greatest for pasture biomass and lowest for CPUE (Figs 4 & 5). For instance, if flow decreased by multiples of 0.9 (10% per year), the time required for mean CPUE to fall below the 5% quantile (90% probability intervals) of historical variation was 16 years, but >30 years for pasture biomass and NDVI. Emergence times were lower for higher quantiles, reflecting the reduced level of confidence that the trend had emerged from background variation. The differences in emergence times among indicators became smaller as the quantiles were increased; for example at a quantile of 40% the indicators had similar emergence times for all trends in flow.

**Figure 5.**
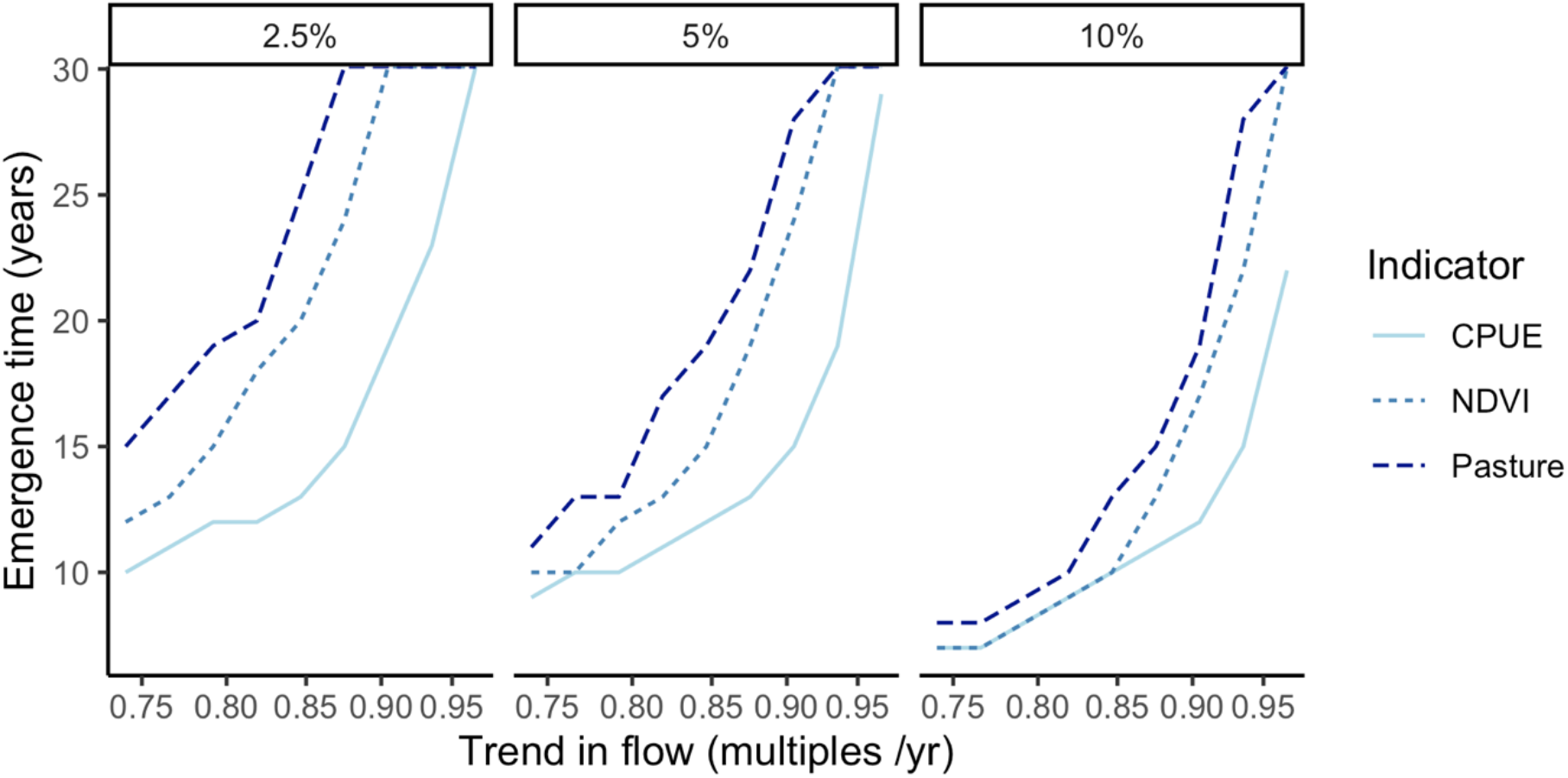
Emergence times for the three indicators, each panel represents a different lower bound: 2.5% = 95% probability intervals, 5% = 90% probability intervals and 10% = 80% probability intervals.

Code and data to repeat the analyses is available at: https://github.com/cbrown5/ecological-condition-latent-model.

## Discussion

We identified relationships among all indicators and flow, and particularly for barramundi catch per unit effort and NDVI. There is high interannual variation in the Mitchell catchment, driven by interannual variation in rainfall. Wet years drive increased vegetation growth and barramundi production, which was reflected in correlations among river flow, NDVI and barramundi catch. Thus, ecological condition, defined here as covariation in the indicators, largely reflected year-to-year variation in flow. This variation underscores the importance of quantifying interconnections between assets via river flow and among ecosystem services. The implication is that change in one variable (flow) can cause changes in multiple processes, and consequently there would be correlated changes in the annual capacity of the system to deliver different ecosystem services.

By defining the ecological condition index as a latent variable we were able to partition out environmental effects shared by all indicators (captured in the ecological condition index) from those that were unique to specific indicators (variation in indicators that is not explained by the condition index). The ability to identify shared trends may be desirable in an indicator, because it facilitates the attribution of changes that affect only specific indicators versus those that impact the overall condition of the ecosystem (Sutherland et al. 2016; Kupschus et al. 2016). Over time there was an increasing trend in barramundi CPUE from the 1990s to 2015. This was driven by high flows 2005-2015 and management changes in the fishery in the late 1990_s_ and early 2000_s_, which reduced fishing effort and put increased restrictions on fishing gear (Streipert et al. 2019). NDVI had a strong correlation with flow and CPUE, but in contrast to CPUE, NDVI did not show as strong a long-term trend, because it was not affected by fishery management changes. Pasture biomass had the weakest correlation with the other indicators, perhaps because it was more strongly driven by rainfall that is local to a region and less by inundation caused by increased river flow.

The typical approach to defining joint indices of multiple ecosystem components has been spatial and thematic aggregation by summing spatially-specific counts or by averaging and then normalising spatially-specific indicators (Maes et al. 2020). The aggregation procedure requires the use of some form(s) of aggregation functions and weights. Our model-based approach to defining a condition index can also be interpreted as a weighted sum of the indicators, where the weights were estimated empirically. Each approach has its own advantages and disadvantages. Our approach requires time-series data, which are not available for many indicators or in some regions (Dvarskas 2018). Expert weights do not require historical data and are simple to compute, but can also be arbitrary. Arbitrary weights are not repeatable across different groups of experts and may mis-represent management priorities (Game et al. 2013). Further, our approach partitioned variation in CPUE caused by flow from changes in management of the fishery. Simply summing weighted CPUE with other indicators would tend to overinflate the influence of barramundi fishery management on ecological condition as a whole. Thus, a key advantage of our approach is that it can partition changes unique to individual indicators, versus changes that are shared across the ecosystem. Identifying redundancy and complementarities among different indicators is an important step in ensuring that monitoring programs can focus on a parsimonious set of indicators that is comprehensive of the ecosystem’s dynamics (Czúcz et al. 2021).

A number of modelling toolboxes have been previously developed to quantify connections among the condition of ecosystems and ecosystem service flows, but none of these directly address the issue of how effectively one can detect trends in indicators. Examples include the soil and water assessment tool (Bieger et al. 2017), the Integrated valuation of ecosystem services tradeoff (INVEST) tool (Guerry et al. 2012) and globally applicable artificial intelligence tools (Martínez-López et al. 2019). These tools have been applied for measuring connected ecosystem services across catchments, in particular regulating ecosystem services relating to water quantity, water quality and sediment erosion (Bieger et al. 2017; Hamel et al. 2017). All share a common approach, based on parameterising models of soil erosion and run-off and then analysing predicted service flows spatially in a geographic information system. However, these toolboxes are designed for making service flow projections, given sufficient information exists to parameterize their models. The utility of our approach is as a performance evaluation tool, which is an understudied aspect of biodiversity indicator development (Rowland et al. 2020; Watermeyer et al. 2021). Our approach, which blends state-space and structural equation modelling, could be extended to model multivariate change in other ecosystems. This type of model warrants further development for assessing ecosystem level change, particularly where there is a need to differentiate the different sources of stochasticity in observed time-series (Auger-Méthé et al. 2021)

The high interannual variation in flow and ecological condition will create challenges for detecting change in the Mitchell catchment. Our estimates of the interannual variation for each indicator were broad and thus our model predicted that emergence times across all indicators at typical confidence bounds (10% or 5%) were >10 years and sometimes >30 years for all but the most extreme trends in flow. These long emergence times mean the indicators will not be useful for managers aiming to use the model to pre-emptively respond to change, because the changes will not be detected on a management relevant timespan (Stevenson et al. 2021). Rather, the indicators will capture long-term low-bandwidth changes in the ecosystem. For comparison, proposed water developments in the Mitchell catchment are expected to cause much smaller reductions in flow, in order of 1-5% per year (Petheram et al. 2018), than most of the trends we explored here. Such water developments however are predicted to impact estuarine ecosystem services (Broadley et al. 2020; Leahy & Robins 2021). Thus, our findings suggest management planning should not rely on indicators of ecological condition. Instead, management should use pressure indicators, that may better pre-empt change in ecological condition (Czúcz et al. 2021). Predictive modelling of the response to ecosystem services to future changes in land-use and water flow is also needed to inform pre-emptive management in the Mitchell catchment (McMahon et al. 2022).

There are some important caveats to consider in our analysis. First, there was missing data in some of the indicator time-series. Missing data in time-series is a common problem in modelling and will reduce the power to detect long-term change (Dvarskas 2018). The latent variable approach enabled interpolating missing data, on the strength of correlated variables (Hui et al. 2015). A simulation study suggested accuracy of predictions for a variable was not sensitive to missing data in other variables. Thus, we suggest our approach could have value more generally for creating indices of ecological condition when there are gaps in time-series.

The analysis of barramundi used a simplified stock assessment model. For many fisheries additional data may be available, such as data on age structure, and this could improve the accuracy of the modelling. Our simple stock assessment did compare well with a more quantitatively complex stock assessment, with the stock status estimates being similar (Streipert et al. 2019) – this is the consequence of both assessment models being driven primarily by trends in CPUE and the assumption of an depletion of 0.2 B_0_ in 1990. More importantly, the availability of barramundi to estuarine fisheries, as well as productivity, is also likely to be affected by flow (Robins et al. 2005). Future analyses could incorporate this effect if more data were available to separate catchability from productivity effects.

An important caveat is that our indicators are an incomplete representation of the processes supporting capacity to supply ecosystem services in the Mitchell catchment. We focussed on assets that support commercial fish harvest and commercial livestock grazing, because there were sufficient time-series data for these indicators to assess emergence times. However, Indigenous uses of ecosystems are also an important component of the ecosystem services in Northern Australian catchments (Scheepers & Jackson 2012; Jackson et al. 2012, 2014), which may be particularly sensitive to development (Stoeckl et al. 2013). Our modelling did not include condition directly relating to Indigenous services, due to lack of time-series data. It is likely that our conclusions regarding emergence time would also apply to those assets supporting traditional uses, such as water holes, given the interconnections in this catchment. Future modelling studies could integrate Indigenous knowledge to enhance the predictive power of ecosystem service models. For example, Indigenous knowledge on the spatial distribution of barramundi (Scheepers & Jackson 2012) could be incorporated into ecosystem service models (e.g. Guerry et al. 2012). This would enhance our ability to predict change in ecosystem services, as well as quantify their value in a broader range cultural contexts.

## Conclusion

We analyzed change in indicators for interconnected ecosystem assets and found a high level of variation. This high level of variation means long time-series are required to detect change in the capacity of the ecosystem to supply ecosystem services. Our quantification of timescales to detect change provides important context for policy-makers when they are considering indicators (Stevenson et al. 2021). For instance, we urge caution when using indicators like the ones analysed here in cost-benefit analysis when evaluating water resource development proposals. Economic variables may respond rapidly and positively to development (Stevenson et al. 2021), but the consequent degradation of ecological condition and ecosystem’s capacity to supply ecosystem services may take much longer to be noticed. These temporal differences in response times may not be adequately captured in cost benefit analysis, which may lead to decisions that are biased towards development opportunities that bring short-term benefits but have long-term environmental costs.

## Notes

https://github.com/cbrown5/ecological-condition-latent-model

### Competing Interest Statement

The authors have declared no competing interest.

### Summary of Updates

Reframe the manuscript to be about performance testing of indicators.

