## Supplementary material for "Ecosystem services in connected catchment to coast ecosystems: monitoring to detect emerging trends"

Supplemental material for: Ecosystem services in connected catchment to coast ecosystems: monitoring to detect emerging trends

Christopher J. Brown* [1], Chantal Saint Ange [2], Rod M. Connolly [1], Syezlin Hasan [2], Sue Jackson [2], Joseph M. McMahon [2], James C. R. Smart [2]

1. Coastal and Marine Research Centre, Australian Rivers Institute, School of Environment and Science, Griffith University, Gold Coast, QLD 4222, Australia

2. Australian Rivers Institute, School of Environment and Science, Griffith University, Nathan, QLD, Australia

**Supplemental methods**

**Table S1** Data on the variables and indicators, how it relates to the account and its source.

| Indicator | Source | Years | References |
|---|---|---|---|
| Average streamflow (m^3/s) | CSIRO | 1990-2015 (wet-season years) | Hughes et al. (2017)<br>https://nawra-river.shinyapps.io/river/ |
| Barramundi catch (Tonnes) | Fisheries Queensland | 1990-2017 | State of Queensland Department of Agriculture Fisheries and Forestry (2018)<br>https://qfish.fisheries.qld.gov.au/ |
| Effort (Days) | Fisheries Queensland | 1990-2017 | State of Queensland Department of Agriculture Fisheries and Forestry (2018)<br>https://qfish.fisheries.qld.gov.au/ |
| Normalized difference vegetation index (NDVI) | Bureau of Meteorology | 1992-2019 | http://www.bom.gov.au/climate/austmaps/about-ndvi-maps.shtml |

| | | | |
|---|---|---|---|
| Pasture Biomass – Total standing dry matter (kg DM/ha) | Long Paddock | 1990-2015 | State of Queensland Department of Environment and Science (2021) https://www.longpaddock.qld.gov.au/aussiegrass/ |

*Prior choices*

We constrained beta_nu to positive values by using an exponential prior (Table S1). Beta_u, beta_pasture and beta_ndvi were also scale invariant with beta_nu ($\beta^{\nu}$, i.e. high values of beta_nu resulted in lower values of the regressions of the indicators on the ecological condition nu). Therefore, we set their priors so that they all had equal scales (Table S1). Priors for the state-space model were chosen to constrain the model to estimation with an expected range for population parameters, interannual variation in biomass and CPUE (e.g. Millar and Meyer, 2000).

**Table S2** Prior choices

| Parameter | Prior options | Reasoning |
|---|---|---|
| Beta [pasture biomass and NDVI] | Normal(0, 1) | The normal prior allows either a negative or positive relationship with ecological condition (and therefore flow). Setting SD = 1 meant these parameters had similar scale to beta_nu. |
| Beta u [fish stock process errors] | Normal(0, 0.2) | Normal prior for the lognormal (multiplicative) fish stock process errors. The chosen SD means the 90% quantiles for this prior range over a multiplicative effect of 0.73-1.40. This ensures model identifiability and efficient convergence. |
| a [pasture biomass and NDVI] | Normal(0, 10) | Broad, very weakly informed prior (note CPUE and NDVI were scaled to unit variance). |
| Beta nu ($\beta^{\nu}$) | Exponential(1) | The exponential prior constrained the sign to be positive and aided identification of parameters. A rate of 1 meant the scale of this parameter was the same as the scale for the other beta parameters. |
| Ln(K) | Normal(7.91, 0.83) | Chosen assuming biomass in 1990 (year one) was 20% of unfished and that catch in year 1 was 10% of biomass in year 1. Assuming a CV of 1. |

| | | |
|---|---|---|
| ln(r) | Normal((-1.24, 0.29) | Chosen to have mean of 0.3 and CV of 0.3. Based on barramundi life-history traits. |
| q | Uniform(0.001, 0.15) | Calculated range based on scale of catch values and assumed initial depletion level (Millar and Meyer, 2000). |
| Sigma (ndvi, inun) | Exponential(0.1) | Weakly informative prior on observation error SD that shrinks estimates to zero (McElreath, 2020). |
| Sigma (cpue) | Exponential(2) | Moderately informative prior on observation error SD that shrinks estimates to zero (McElreath, 2020). Using a slightly greater shrinkage on CPUE improved convergence. |
| Sigma u | Exponential(14) | Chosen so prior probability of process error would deviate from 5% was <0.95 |

*Algorithm settings*

The model was estimated with the STAN programming language, implemented from the R program with the rstan package (Team, 2020). Algorithm convergence was achieved with 5,000 iterations across 3 chains and a thinning rate of 5. Convergence was confirmed with Rhat statistics < 1.01 and via visual diagnostics.

*Sensitivity to catchability change and initial biomass*

Our main analyses assumed catchability was constant over time and that the initial biomass of barramundi was a fixed fraction of the carrying capacity. We tested the effects of these assumptions on parameter estimates and model fits by varying the two parameters in sensitivity analyses.

First, it is possible that gear changes in the fishery have caused increases in effective fishing effort over time. We therefore re-fit the model assuming a 1% increase in catchability per-annum.

Second, we assumed that initial biomass was 20% of the carrying capacity, based on the latest stock assessment (Streipert et al., 2019). However, the stock assessment was for the northern and southern, gulf, whereas the Mitchell catchment, in the southern Gulf, may have been more lightly fished than the northern Gulf in the 1990s. Therefore, we refit our model with initial biomasses that were 50% and 80% of carrying capacity.

Finally, we crossed the above two comparisons, running both catchability scenarios for all three initial biomass ratios.

We analysed results by comparing parameter estimates from the models and also the Bayesian r-squared (Gelman et al., 2019).

*Simulation study*

We ran a simulation study where we varied the amount of missing data for the NDVI dataseries and compared estimated parameters to known simulated values.

We first fit an ARIMA model to the standardized flow data, using the R package ‘forecast’ (Hyndman and Khandakar, 2008). The optimal model was selected using the AIC and was found to have order 6, 0, 3. We then simulated 50 by 26 year flow series using the fitted ARIMA model. Then we simulated observations of NDVI, pasture biomass and CPUE using the median parameter values from the fitted model. We then fit the model to the simulated data-series and recorded median parameter values with 50% and 95% credible intervals. We repeated this process where the first 0, 5, 11 and 15 years of the NDVI data were missing in when we refit the model to the simulated data.

We then compared how the model estimated parameter values compared to the ‘true’ value used to set-up the simulations, and how well the fitted model’s predicted the simulated data they were fitted to. We compared the values with coverage statistics and the Bayesian $R^2$ between the simulated data-series and each model’s predicted values. The Bayesian $R^2$ was defined as the distribution of $R^2$ values calculated across the posterior distribution of predictions for each variables.

*Direct and total effects of condition and flow on NDVI, pasture biomass and barramundi surplus production*

For pasture biomass and NDVI, the expected difference in values from year *a* to year *b* can be derived:

$$\mathrm{y}_a - \mathrm{y}_b = \beta^u \beta^v ln\left({F_a}/{F_b}\right) \qquad \text{(eqn S1)}$$

For barramundi first define surplus production, SP, as:

$$\mathrm{SP}_t = B_t - B_{t-1} + C_t \qquad \text{(eqn S2)}$$

Which gives:

$$\mathrm{SP}_t = rB_{t-1}\left(1 - \frac{B_{t-1}}{K}\right)u_t \qquad \text{(eqn S3)}$$

Expanding the term for $u_t$ gives us:

$$\mathrm{SP}_t = rB_{t-1}\left(1 - \frac{B_{t-1}}{K}\right)exp\left(\beta^u\left(\beta^v \ln\left(F_t\right) + \epsilon_t^\gamma\right) + \epsilon_t^u\right) \qquad \text{(eqn S4)}$$

The error term for ecological condition ($\epsilon_t^v$) is distributed as a standard normal variate, so the retransformation bias induced in the mean of $SP_t$ is equal to a factor of exp(0.5) = 1.65. The error term for the process errors ($\epsilon_t^u$) will induce a bias in the mean of $\exp\left(0.5\sigma^u\right)$ (Stow et al., 2006). In the main results we present the posterior for $SP_t$ with quantiles which are unbiased by non-linear transformation, so we do not consider the retransformation bias further.

The relative effect of a change in flow or condition on surplus production from year *a* to year *b* can then be derived:

$$\mathrm{SP}_a/\mathrm{SP}_b = {F_a}/{F_b}\, exp(\beta^u\beta^v) \qquad \text{(eqn S5)}$$

**Supplemental results**

*Model verification*

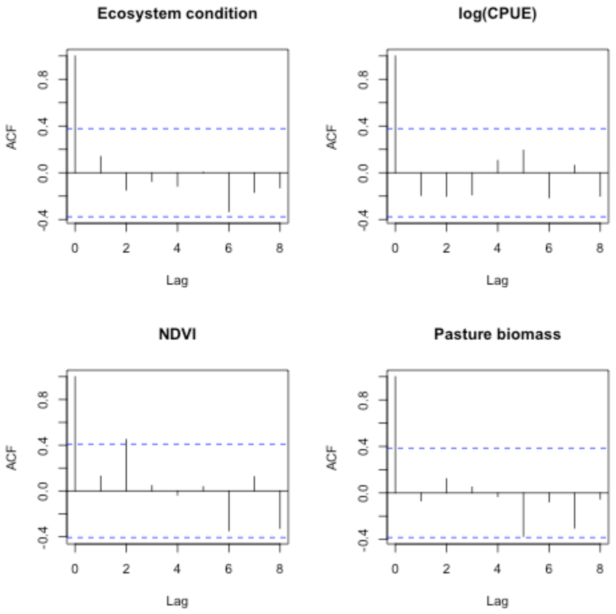


**Figure S1** Autocorrelation in residuals of key variables at lags of 0 to 8 years. Blue dotted line indicates values that are statistically significant at the 5% probability level.

*Sensitivity analyses*

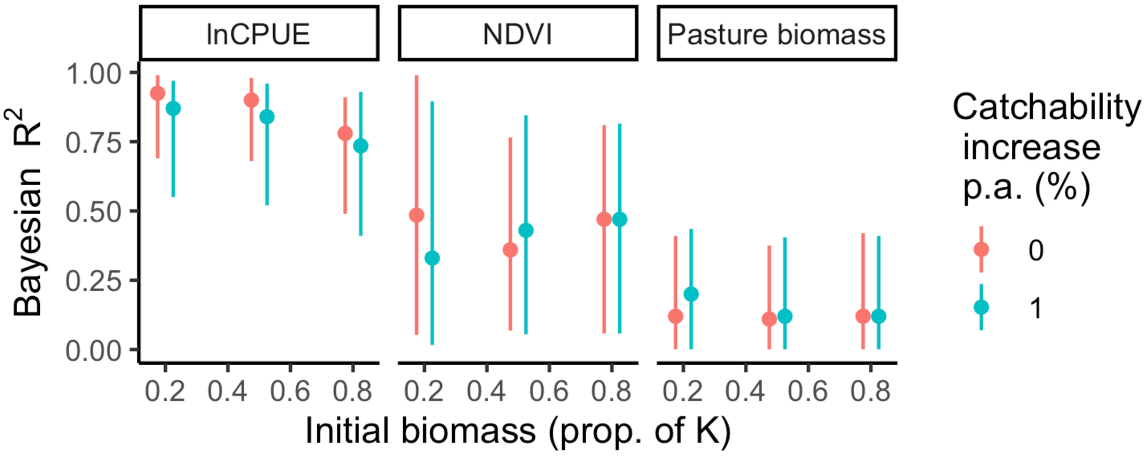


**Figure S2** Bayesian $R^2$ values for the main results and sensitivity analyses where the rate of catchability increase and the initial biomass ratio were varied. Points show medians, bars show 95% CIs.

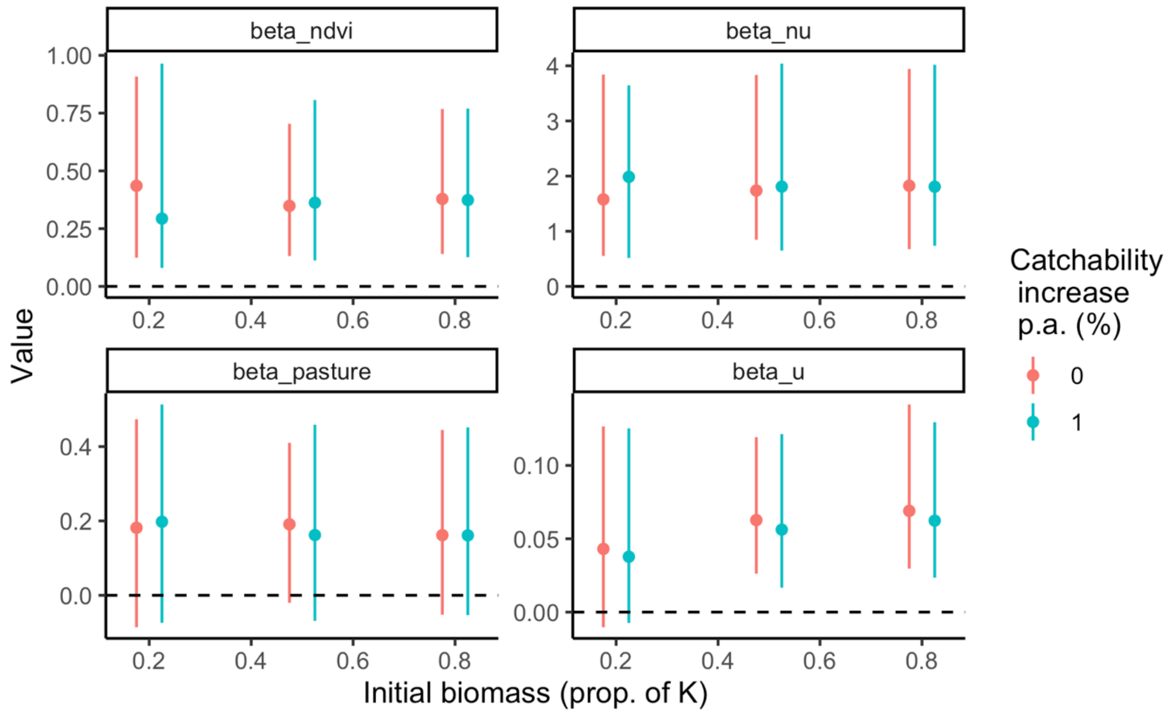


**Figure S3** Beta parameters from the sensitivity analyses where the rate of catchability increase and the initial biomass ratio were varied. where the model was fitted to 50 replicate randomized time-series of flow, NDVI and inundation. Points show medians, bars show 95% CIs.

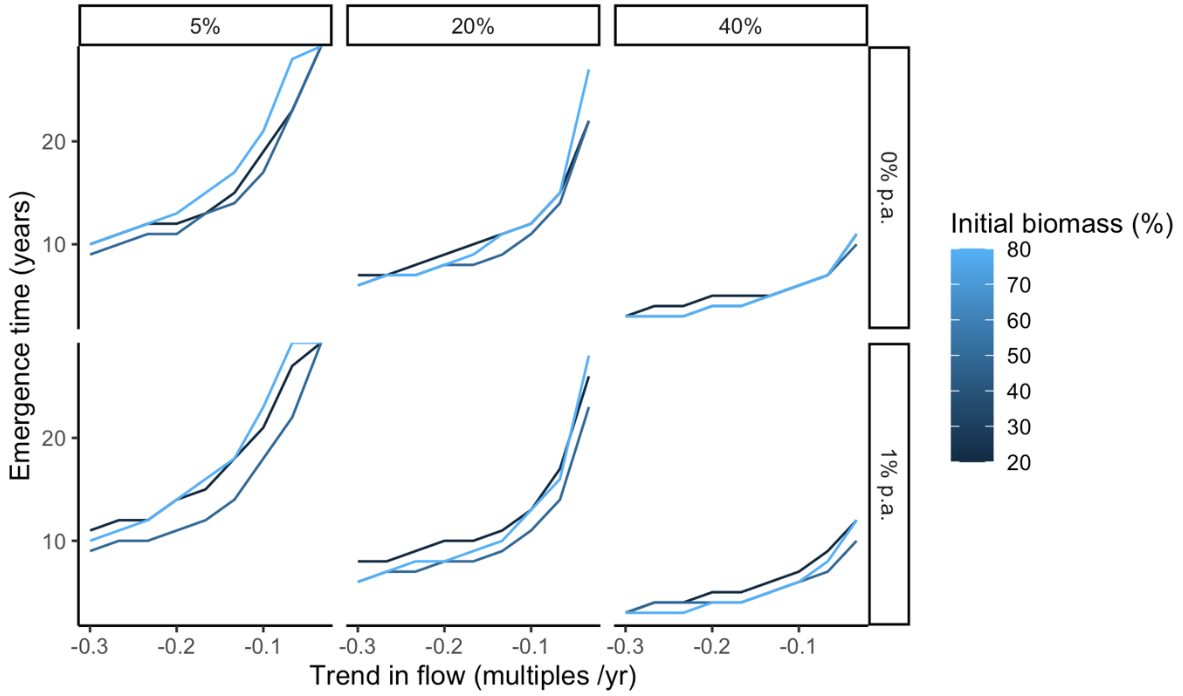


**Figure S4** Emergence times for CPUE from the sensitivity analyses where the rate of catchability increase and the initial biomass ratio were varied. Columns show different quantiles for emergence times, rows show different % increase in catchability per annum, colours show the assumed biomass in 1990, as a percent of carrying capacity.

*Simulation study results*

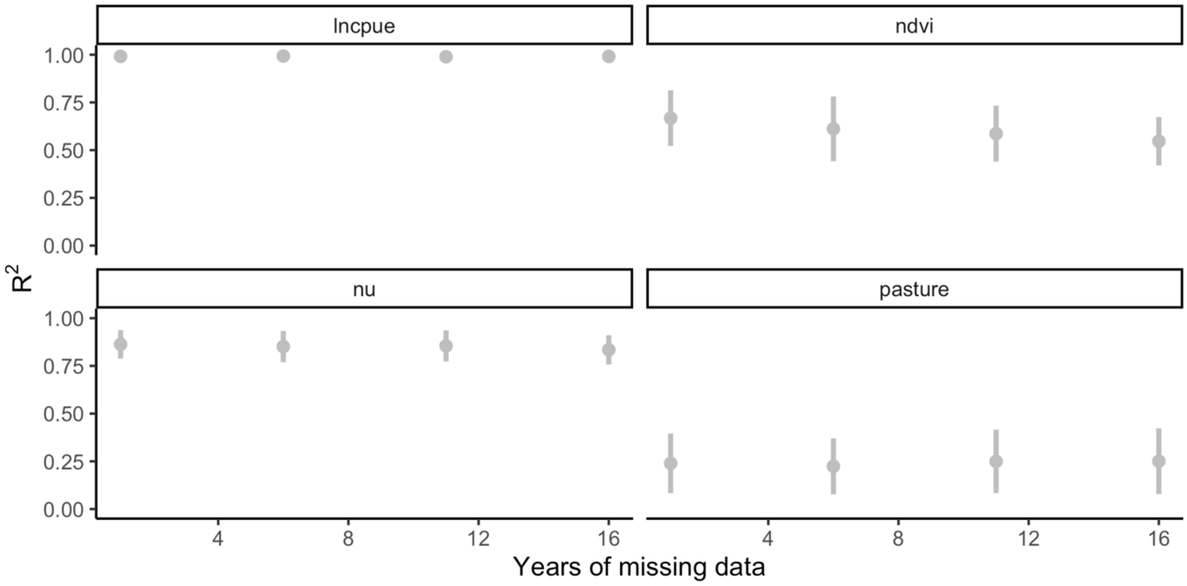


**Figure S5** Results of simulation test, showing correlation between simulated data values and data values predicted by fitted models. Simulations were performed with the first 0, 5 and 10 years of inundation data missing. Points show mean $R^2$ values, bars show 1 standard deviation above and below the mean. Panels show each variable, including 'nu' the latent ecological condition.

**Table S3** Proportional coverage of 95% and 50% CIs of the true value from the simulation study where the model was fitted to 50 replicate randomized time-series of flow, NDVI and pasture biomass. Simulations were performed with the first 0, 5 and 10, 15 years of NDVI data missing.

| **Parameter** | **Years missing** | **num valid simulations** | **Coverage at 95%** | **Coverage at 50%** |
|---|---|---|---|---|
| a_ndvi | 0 | 41 | 1.00 | 0.00 |
| a_ndvi | 5 | 40 | 1.00 | 0.48 |
| a_ndvi | 10 | 40 | 1.00 | 1.00 |
| a_ndvi | 15 | 44 | 1.00 | 0.52 |
| a_pasture | 0 | 45 | 1.00 | 0.98 |
| a_pasture | 5 | 39 | 1.00 | 0.92 |
| a_pasture | 10 | 41 | 1.00 | 0.93 |
| a_pasture | 15 | 43 | 1.00 | 0.98 |
| beta_ndvi | 0 | 45 | 1.00 | 0.27 |

| | | | | |
|---|---|---|---|---|
| beta_ndvi | 5 | 36 | 1.00 | 0.42 |
| beta_ndvi | 10 | 39 | 1.00 | 0.33 |
| beta_ndvi | 15 | 45 | 1.00 | 0.51 |
| beta_nu | 0 | 44 | 1.00 | 0.16 |
| beta_nu | 5 | 40 | 1.00 | 0.10 |
| beta_nu | 10 | 38 | 0.97 | 0.08 |
| beta_nu | 15 | 42 | 0.98 | 0.02 |
| beta_pasture | 0 | 43 | 0.98 | 0.67 |
| beta_pasture | 5 | 36 | 0.97 | 0.44 |
| beta_pasture | 10 | 39 | 0.97 | 0.54 |
| beta_pasture | 15 | 41 | 0.98 | 0.44 |
| beta_u | 0 | 44 | 1.00 | 0.61 |
| beta_u | 5 | 36 | 1.00 | 0.47 |
| beta_u | 10 | 35 | 0.97 | 0.26 |
| beta_u | 15 | 38 | 0.97 | 0.24 |
| lnK | 0 | 41 | 1.00 | 1.00 |
| lnK | 5 | 36 | 1.00 | 0.94 |
| lnK | 10 | 36 | 1.00 | 0.92 |
| lnK | 15 | 38 | 1.00 | 0.95 |
| lnr | 0 | 45 | 1.00 | 0.96 |
| lnr | 5 | 38 | 1.00 | 0.87 |
| lnr | 10 | 38 | 1.00 | 0.82 |
| lnr | 15 | 37 | 1.00 | 0.89 |
| q | 0 | 39 | 1.00 | 0.90 |
| q | 5 | 33 | 1.00 | 0.82 |
| q | 10 | 35 | 1.00 | 0.91 |
| q | 15 | 38 | 1.00 | 0.89 |
| sigma_cpue | 0 | 33 | 1.00 | 0.64 |
| sigma_cpue | 5 | 28 | 1.00 | 0.43 |
| sigma_cpue | 10 | 32 | 1.00 | 0.69 |

| | | | | |
|---|---|---|---|---|
| sigma_cpue | 15 | 36 | 1.00 | 0.42 |
| sigma_ndvi | 0 | 38 | 1.00 | 0.45 |
| sigma_ndvi | 5 | 30 | 1.00 | 0.60 |
| sigma_ndvi | 10 | 42 | 1.00 | 0.64 |
| sigma_ndvi | 15 | 45 | 1.00 | 0.80 |
| sigma_pasture | 0 | 44 | 1.00 | 0.00 |
| sigma_pasture | 5 | 40 | 1.00 | 0.00 |
| sigma_pasture | 10 | 39 | 1.00 | 0.00 |
| sigma_pasture | 15 | 43 | 1.00 | 0.00 |

State of Queensland Department of Agriculture Fisheries and Forestry, 2018. QFish data cube.

State of Queensland Department of Environment and Science, 2021. The Long Paddock.

Stow, C.A., Reckhow, K.H., Qian, S.S., 2006. A Bayesian approach to retransformation bias in transformed regression. Ecology 87, 1472–1477.

Streipert, S., Filar, J., Robins, J., Whybird, O., 2019. Stock assessment of the barramundi (Lates calcarifer) fishery in Queensland, Australia. May 2019. Queensland Government.

Team, S., 2020. RStan: the R interface to Stan. R package version 2.21. 1.
